# Process proteomics of beer reveals a dynamic proteome with extensive modifications

**DOI:** 10.1101/234252

**Authors:** Benjamin L. Schulz, Toan K. Phung, Michele Bruschi, Agnieszka Janusz, Jeff Stewart, John Mehan, Peter Healy, Amanda S. Nouwens, Glen P. Fox, Claudia E. Vickers

**Affiliations:** School of Chemistry and Molecular Biosciences, The University of Queensland, Brisbane, Queensland 4072, Australia; ARC Training Centre for Biopharmaceutical Innovation, Australian Institute of Bioengineering and Nanotechnology, The University of Queensland, Brisbane, Queensland 4072, Australia; Australian Institute of Bioengineering and Nanotechnology, The University of Queensland, Brisbane, Queensland 4072, Australia; Cargill Malt, Adelaide, South Australia 5069 Australia; Lion, Sydney, New South Wales 2127, Australia; Lion, Brisbane, Queensland 4064, Australia; Queensland Alliance for Agriculture and Food Innovation, The University of Queensland, Brisbane, Queensland 4072, Australia

**Keywords:** beer, malt, wort, SWATH, LC-MS/MS, relative quantification, post-translational modifications

## Abstract

Modern beer production is a complex industrial process. However, some of its biochemical details remain unclear. Using mass spectrometry proteomics, we have performed a global untargeted analysis of the proteins present across time during nano-scale beer production. Samples included sweet wort produced by a high temperature infusion mash, hopped wort, and bright beer. This analysis identified over 200 unique proteins from barley and yeast, emphasizing the complexity of the process and product. We then used data independent SWATH-MS to quantitatively compare the relative abundance of these proteins throughout the process. This identified large and significant changes in the proteome at each process step. These changes described enrichment of proteins by their biophysical properties, and identified the appearance of dominant yeast proteins during fermentation. Altered levels of malt modification also quantitatively changed the proteomes throughout the process. Detailed inspection of the proteomic data revealed that many proteins were modified by protease digestion, glycation, or oxidation during the processing steps. This work demonstrates the opportunities offered by modern mass spectrometry proteomics in understanding the ancient process of beer production.

## Introduction

The art of brewing alcoholic beverages has existed for almost the entire length of human civilization as evidenced by records of the Sumerian beer making process recorded on clay tablets from ancient Mesopotamia [1, 2]. While clear advances have been made in beer production since the first bowl of fermenting starchy grain products, aspects of the complex and sequential biochemistry underlying much of the brewing process remain poorly understood.

The process of beer brewing begins with barley grains. These are malted to allow partial germination, triggering expression of key enzymes including glucanases, proteases, α-amylase, β-amylase, and limit dextrinase [2]. This results in degradation of the seed cell walls and protein matrix (hordeins and glutelins) in the endosperm but leaves the starch relatively intact. The germinated grains are then dried and milled. Starch, proteins, and other molecules are solubilized during mashing, a controlled hot water infusion. During mashing, solubilized enzymes degrade starch to fermentable oligomaltose sugars [3–5], and digest proteins to produce peptides and free amino acids. Fermentable sugars and free amino acids are required for efficient yeast growth during fermentation [3, 6]. After the mash, the liquid sweet wort is removed from the spent grain, hops are added for bitterness and aroma, and the wort is boiled. After boiling, the cooled hopped wort is inoculated with yeast, and fermentation proceeds to produce bright beer. Typically this bright beer is then filtered, carbonated, packaged, and sold [3].

Many proteins originating from the barley grain and the yeast are present in beer [7–10], and these have been reported to affect the quality of the final product. For example, barley proteins lipid transfer protein LTP1, protein Z, peroxidase, trypsin inhibitors, and α-amylase can affect the colour, viscosity, and texture of beer [2, 11–14]. Proteins present in beer are also often modified by glycation, due to the high concentrations of reducing sugars in wort [14]. Glycation is common in cooked food, where heat can catalyze non-enzymatic glycosylation of proteins, and can influence the taste and properties of the food [12]. It has been suggested that glycation of protein Z, LTP1, and LTP2 can affect beer foam stability and flavor [15]. Despite the importance of these complexities to beer quality, the beer proteome remains poorly understood.

Here, we used SWATH-MS to investigate the proteomic complexity and dynamics of the beer making process, using nanobrewery-scale production of beer made from Commander malt and fermented with yeast used for the iconic Australian Tooheys New lager.

## Methods

### Malt, grist and wort production

Commander malt, at high (45) and low (34) Kolbach index (KI) levels, was supplied by Cargill Malt (Adelaide, Australia). Coarse and fine grists were prepared in a Buhler Miag disc mill at 1.0 mm and 0.2 mm, respectively. Mashing, with four laboratory replicates, used 50 g malt in 365 mL hot liquor at 65°C with constant stirring for 60 min, cooled to room temperature and made up to 450 g prior to filtration. The mash was filtered through Whatman No 597 ½ paper with the first 100 mL returned. To obtain sweet wort samples for proteomic analysis, 6 mL was taken after the return of the first 100 mL. Approximately 340 mL of wort was used for subsequent analysis. Specific gravity was measured with a DMA 45 density meter (Anton Paar, Germany). Boiling was carried out in 500 mL Schott bottles for 30 min. Pelleted hops were obtained from Lion (Brisbane, Australia), powdered in a mortar and pestle, then added at 10 g/400 mL during the boil step. After the wort was cooled to room temperature, 6 mL was sampled to obtain final hopped wort samples for proteomics analysis. Samples were stored at -20°C until further analysis.

### Fermentation

Fermentation of 250 mL of each wort inoculated with Tooheys New lager yeast from Lion was performed at 20°C in 250 mL glass measuring cylinders stoppered with foam and covered with aluminium foil. Wort was inoculated with 10 × 10^6^ cells/mL at 14°C and transferred to a static 20°C incubator. Fermentation was tracked by measuring mass loss due to CO2 volatilization over time [16] and fermentation was deemed complete when mass stabilized. Following fermentation, beers were clarified by centrifugation to remove yeast and haze, cold-stored for 24 h to stabilize, then recentrifuged to yield bright beer. Samples were taken for proteomics and ethanol analysis.

### Beer analyses

Ethanol concentration was measured by ion-exclusion chromatography as described previously [17] using an Agilent 1200 HPLC system and Agilent Hi-Plex H column (300 × 7.7 mm, PL1170-6830) with guard column (SecurityGuard Carbo-H, Phenomenex PN: AJO-4490). Analytes were eluted isocratically with 4 mM H_2_SO_4_ at 0.6 mL/min at 65°C. Ethanol was monitored using a refractive index detector (Agilent RID, G1362A). Total protein concentration was determined using a Bradford assay. Bright beer colour was measured spectrophotometrically by absorbance at 430 nm at room temperature. Foam stability was tested using a Rudin unit. Glassware was acid-washed and dried prior to use. Bright beer was shaken vigorously to degas and left at room temperature for 12 h to equilibrate. The temperature was measured immediately prior to Rudin analysis. Beer (300 mL) was transferred to the Rudin cylinder and CO2 bubbled through. Foam drainage was timed between checkpoints using a stopwatch.

### Sample preparation for mass spectrometry

Independent triplicates of sampled sweet wort, hopped wort, or bright beer were stored at -20°C until analysis. Proteins in 1 mL wort or beer were precipitated by addition of 100 μL of 4 mg/mL sodium deoxycholate in 100% (w/v) trichloroacetic acid (TCA), incubation at 0°C for 30 min, and centrifugation at 18,000 rcf for 10 min. The pellet was resuspended in 1 mL ice cold acetone, vortexed, incubated at 0°C for 15 min, and centrifuged at 18,000 rcf for 10 min. The pellet was air-dried, and proteins were reduced/alkylated by resuspension in 100 μL of 8 M urea, 50 mM ammonium bicarbonate, and 5 mM dithiothreitol and incubation at 56°C for 30 min, followed by addition of iodoacetamide to a final concentration of 25 mM and incubation at room temperature in the dark for 30 min, and addition of dithiothreitol to a final additional concentration of 5 mM. Sample containing 10 μg protein, as determined by 2D quant, was diluted with 50 mM ammonium bicarbonate to a final concentration of 2 M urea. 100 ng trypsin (proteomics grade, Promega) was added and proteins were digested at 37°C for 16 h.

### Mass Spectrometry

Peptides were desalted using C18 ZipTips (Millipore) and analysed by LC-ESI-MS/MS using a Prominence nanoLC system (Shimadzu) and TripleTof 5600 mass spectrometer with a Nanospray III interface (SCIEX) essentially as described [18, 19]. Approximately 0.5-2 μg peptides were desalted on an Agilent C18 trap (300 Å pore size, 5 μm particle size, 0.3 mm i.d. × 5 mm) at a flow rate of 30 μl/min for 3 min, and then separated on a Vydac EVEREST reversed-phase C18 HPLC column (300 Å pore size, 5 μm particle size, 150 μm i.d. × 150 mm) at a flow rate of 1 μl/min. Peptides were separated with a gradient of 10-60% buffer B over 45 min, with buffer A (1% acetonitrile and 0.1% formic acid) and buffer B (80% acetonitrile with 0.1% formic acid). Gas and voltage setting were adjusted as required. An MS TOF scan from *m/z* of 350-1800 was performed for 0.5 s followed by information dependent acquisition of MS/MS with automated CE selection of the top 20 peptides from *m/z* of 400-1250 for 0.05 s per spectrum. SWATH-MS was performed essentially as described [20] with LC conditions identical to those used in information dependent acquisition, but with an MS-TOF scan from an *m/z* of 350-1800 for 0.05 s followed by high sensitivity information independent acquisition with 26 *m/z* isolation windows with 1 *m/z* window overlap each for 0.1 s across an *m/z* range of 400-1250. Collision energy was automatically assigned by the Analyst software (SCIEX) based on *m/z* window ranges. All samples were analysed by SWATH-MS. One randomly selected sample from each set of triplicates was analysed by information dependent acquisition LC-MS/MS.

### Data analysis

Peptides were identified using ProteinPilot 4.1 (SCIEX), searching the LudwigNR database (downloaded from http://apcf.edu.au as at 27 January 2012; 16,818,973 sequences; 5,891,363,821 residues) with standard settings: Sample type, identification; Cysteine alkylation, iodoacetamide; Instrument, TripleTof 5600; Species, none; ID focus, biological modifications; Enzyme, Trypsin; Search effort, thorough ID. False discovery rate analysis using ProteinPilot was performed. Peptides identified with greater than 99% confidence and with a local false discovery rate of less than 1% were included for further analysis. The ProteinPilot data were used as ion libraries for SWATH analyses. The abundance of ions, peptides, and proteins was measured automatically by PeakView Software 2.1 (SCIEX) as previously described [20]. Statistical analyses of protein abundance from SWATH proteomic data were performed as previously described [21] using MSstats in R [22]. Comparison of peptide relative abundance was performed based on peptide intensities [20]. Aligned sequences were visualized using WebLogo [23].

## Results and Discussion

Commercial malting barley was malted at two modification levels (high and low KI), milled to two grist sizes (fine and coarse), and used to produce sweet wort and hopped wort (Fig. 1). Fermentation was commenced by inoculation with Tooheys New lager yeast, and followed by the rate of mass loss over time (Fig. 2A). All conditions resulted in equivalent fermentation rate and extent (Fig. 2A and B). Bright beer after fermentation contained ~3.8% ethanol, while no ethanol was detected before fermentation (Fig. 2C). Total protein concentration was consistent in sweet wort and hopped wort, but substantially decreased after fermentation in bright beer (Fig. 2D). The colour of the bright beer produced from high KI malt was darker than from low KI malt, but showed no dependence on the grist size (Fig. 2E). Foam stability was broadly consistent between conditions, with a significant but small increase in bright beer produced from high KI malt compared to low KI malt (Fig. 2F).

**Figure 1.**
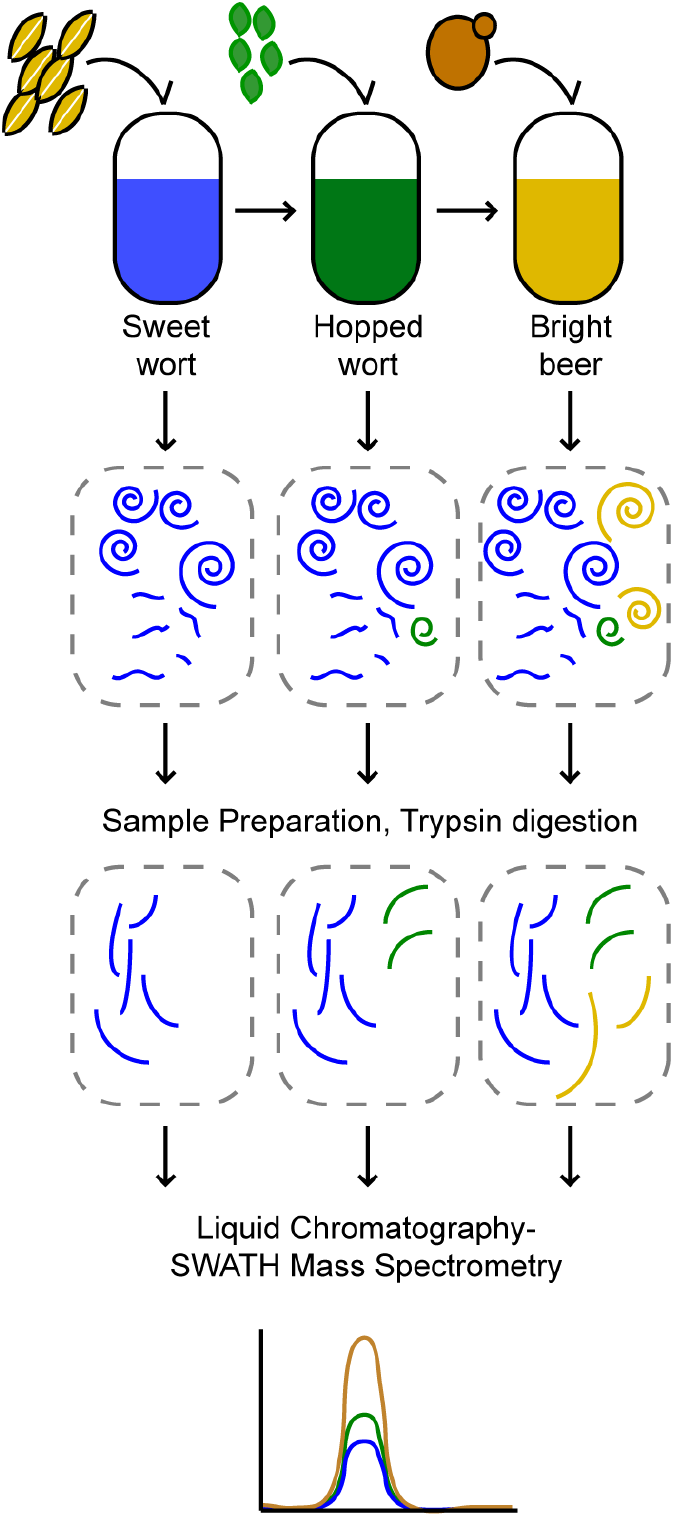
Overview of SWATH-MS proteomics of beer. Malt was prepared at high or low KI, and ground at fine or coarse grist. Beer was produced, with samples taken of sweet wort, hopped wort, and bright beer. Proteins in each fraction were precipitated, denatured, digested to peptides by trypsin, and measured by liquid chromatography-mass spectrometry, with SWATH-MS relative quantification of peptide and protein abundance.

**Figure 2.**
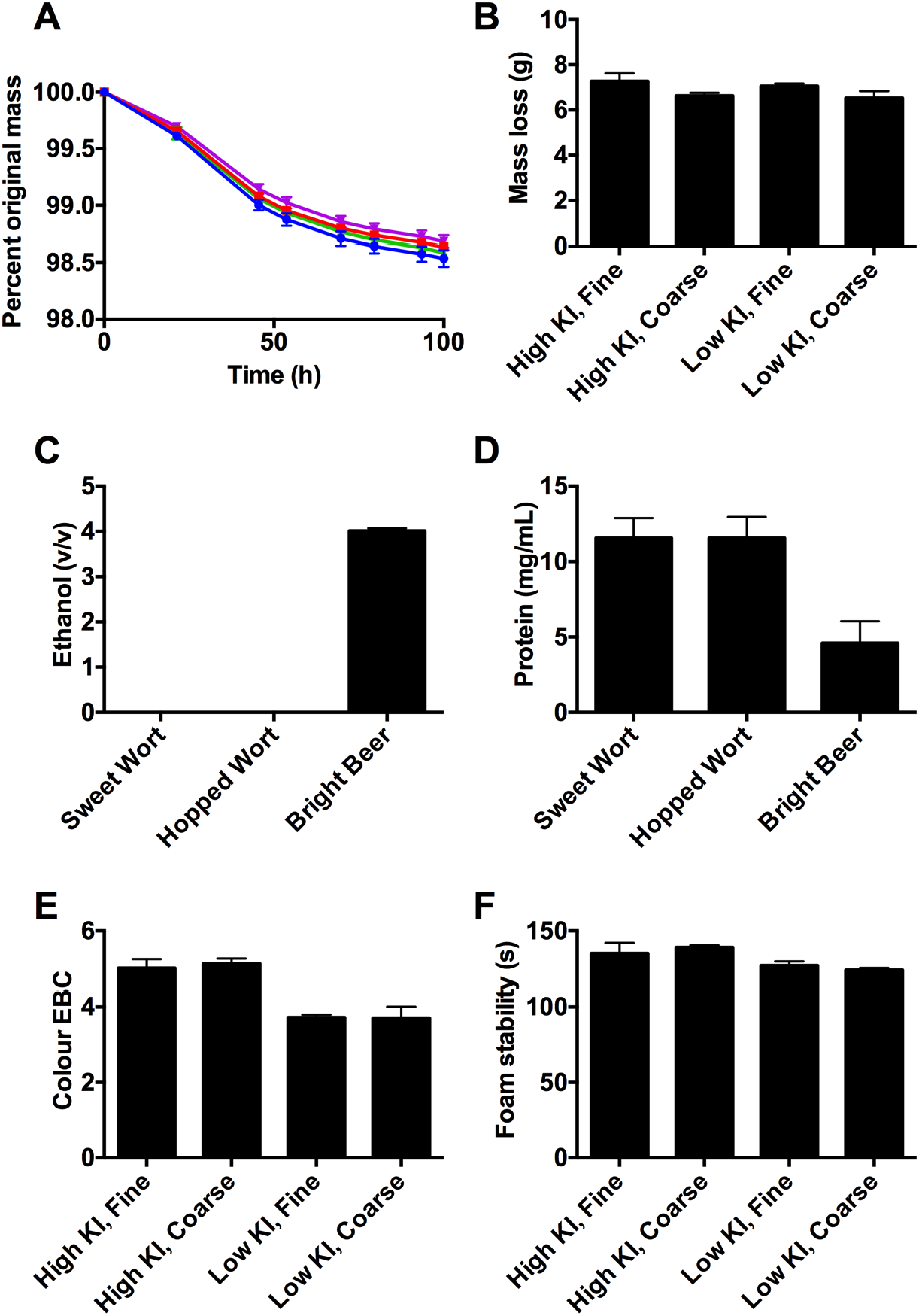
Beer process and product characteristics. (**A**) Fermentation progress monitored by mass loss over time. High KI fine, blue; high KI coarse, red; low KI fine, green; low KI coarse, purple. (**B**) Mass loss over the course of fermentation. (**C**) Ethanol concentrations over the course of the beer production process. (**D**) Total protein concentration over the course of the beer production process. (**E**) Colour. (F) Foam stability. Values are mean, n=4. Error bars are standard deviation.

Beer and wort are complex mixtures of soluble and suspended proteins, carbohydrates, and small organic molecules, and proteomic analysis requires the proteins and peptides be efficiently separated from these other components prior to analysis. Proteins in these samples were separated from other components by TCA precipitation, and were further pre-treated for MS analysis by denaturation, reduction/alkylation of cysteines, digestion to peptides by trypsin, and desalting. Peptides were detected by untargeted LC-MS/MS analysis, and identified by searching peptide MS/MS fragmentation data against the UniProt database of all annotated protein sequences from all species. This identified 210 unique quantifiable proteins from barley and yeast (Supplementary Table S1). Proteins identified were predominantly from barley including lipid transfer proteins, hordeins, and α-amylase inhibitors. Many yeast proteins were identified in bright beer samples, including secreted cell wall enzymes and abundant intracellular metabolic enzymes.

We measured the relative abundance of all proteins in each sample with SWATH-MS. The MS/MS fragmentation spectra and validated list of confidently identified peptides were used to generate a peptide library, which was then used to measure the abundance of each peptide in each sample. The relative abundance of each protein was then compared based on the sum of the intensities of peptides from each protein. To compare the global proteomes of sweet wort, hopped wort, and bright beer, we constructed an unguided clustered heat map based on normalized protein abundance in each sample. This showed the dominant differences between samples were due to the stage in the process (sweet wort, hopped wort, or bright beer), rather than between malting or grist conditions (Fig. 3A). Comparisons of the proteomes of sweet wort, hopped wort, and bright beer for one selected condition (high KI with coarse grist) showed that the largest change in the proteome occurred during fermentation from hopped wort to bright beer (Fig. 3B). The proteins which contributed to these large differences between hopped wort and bright beer were primarily abundant yeast proteins secreted into the wort during fermentation (Table 1).

**Figure 3.**
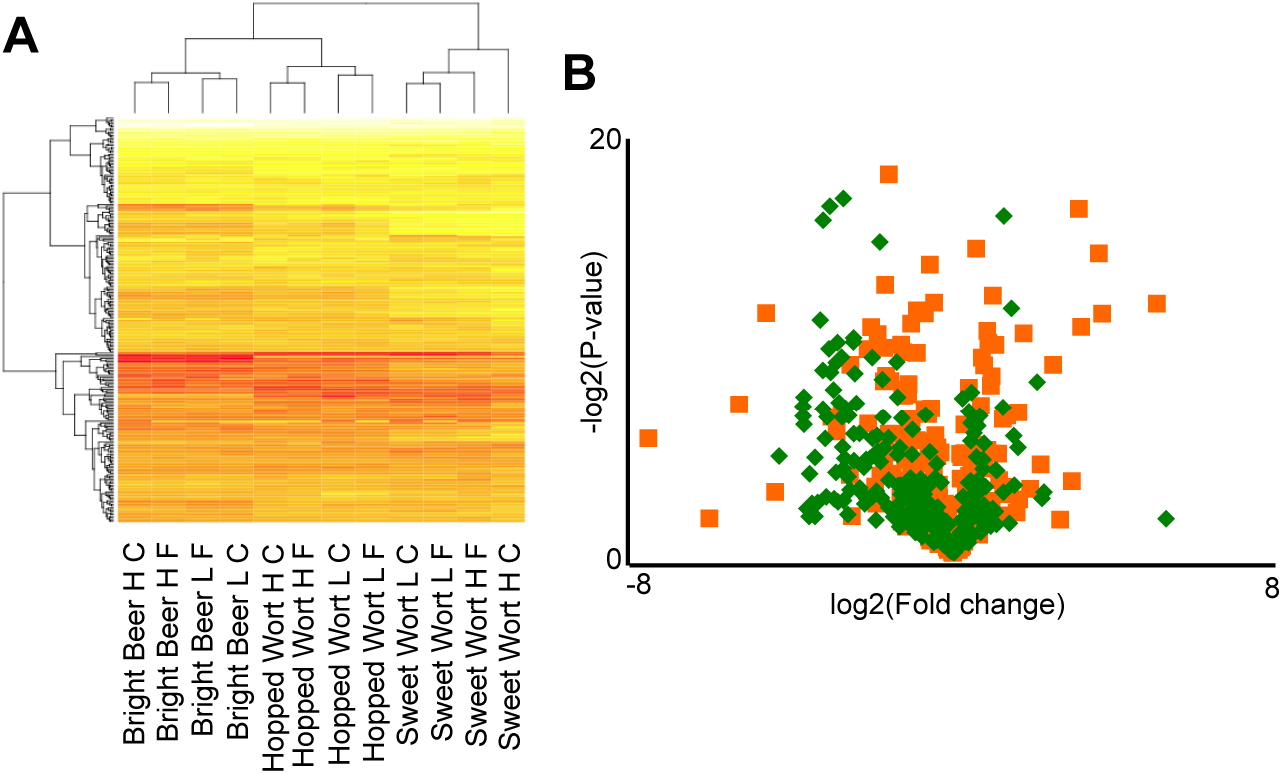
Proteomic comparison of sweet wort, hopped wort, and bright beer. (**A**) Clustered heat map of protein abundance in samples throughout the beer production process, with variation of malting (High KI, H; Low KI, L) and grist parameters (Fine, F; coarse, C). (**B**) Volcano plot comparing the proteomes of hopped wort versus sweet wort (green), and bright beer versus hopped wort (orange).

**Table 1.**
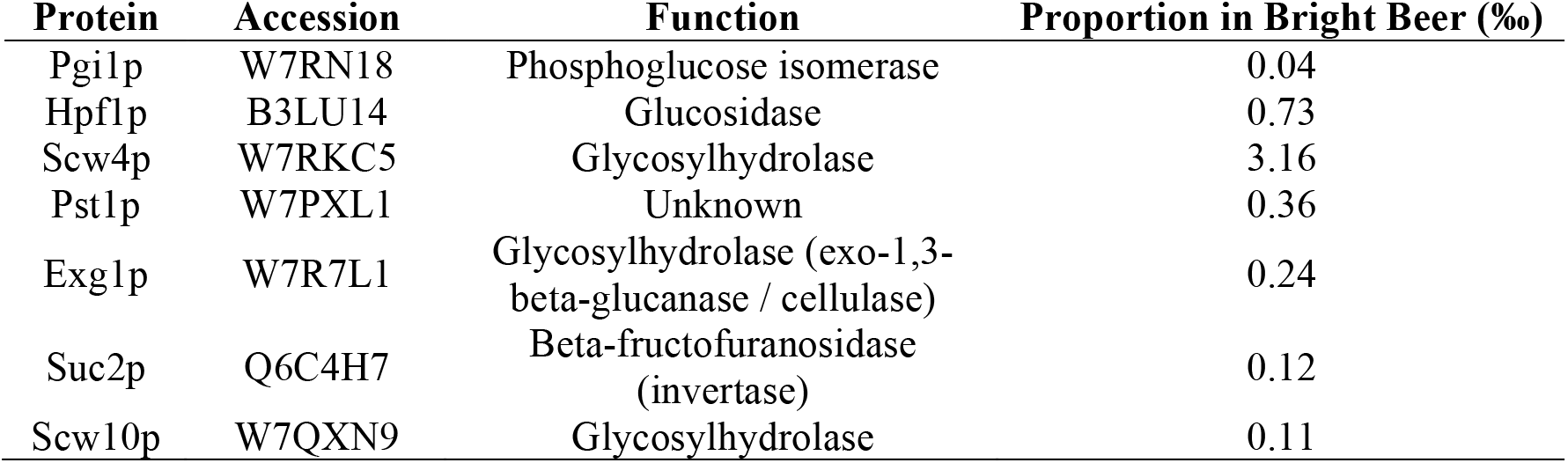
Yeast proteins in bright beer

Our SWATH-MS comparison of sweet wort, hopped wort, and bright beer (Fig. 3, Supplementary Table S2) showed that there were substantial changes to the abundance of many barley proteins at each of these stages during the beer brewing process. To investigate the mechanisms underlying these changes, we compared the biophysical properties of barley proteins with large and significant changes from sweet wort to hopped wort (over the boil), and from hopped wort to bright beer (over fermentation). This showed that high molecular weight proteins were reduced in abundance during the boil (Fig. 4A), while hydrophobic proteins with high grand average hydrophobicity (GRAVY) scores were reduced in abundance during fermentation (Fig. 4H). Other protein biophysical parameters tested showed no significant differences (Fig. 4). These data are consistent with large proteins showing particular sensitivity to thermal unfolding and aggregation during the boil, and with hydrophobic proteins showing sensitivity to organic solvent concentrations from increased ethanol produced during fermentation.

**Figure 4.**
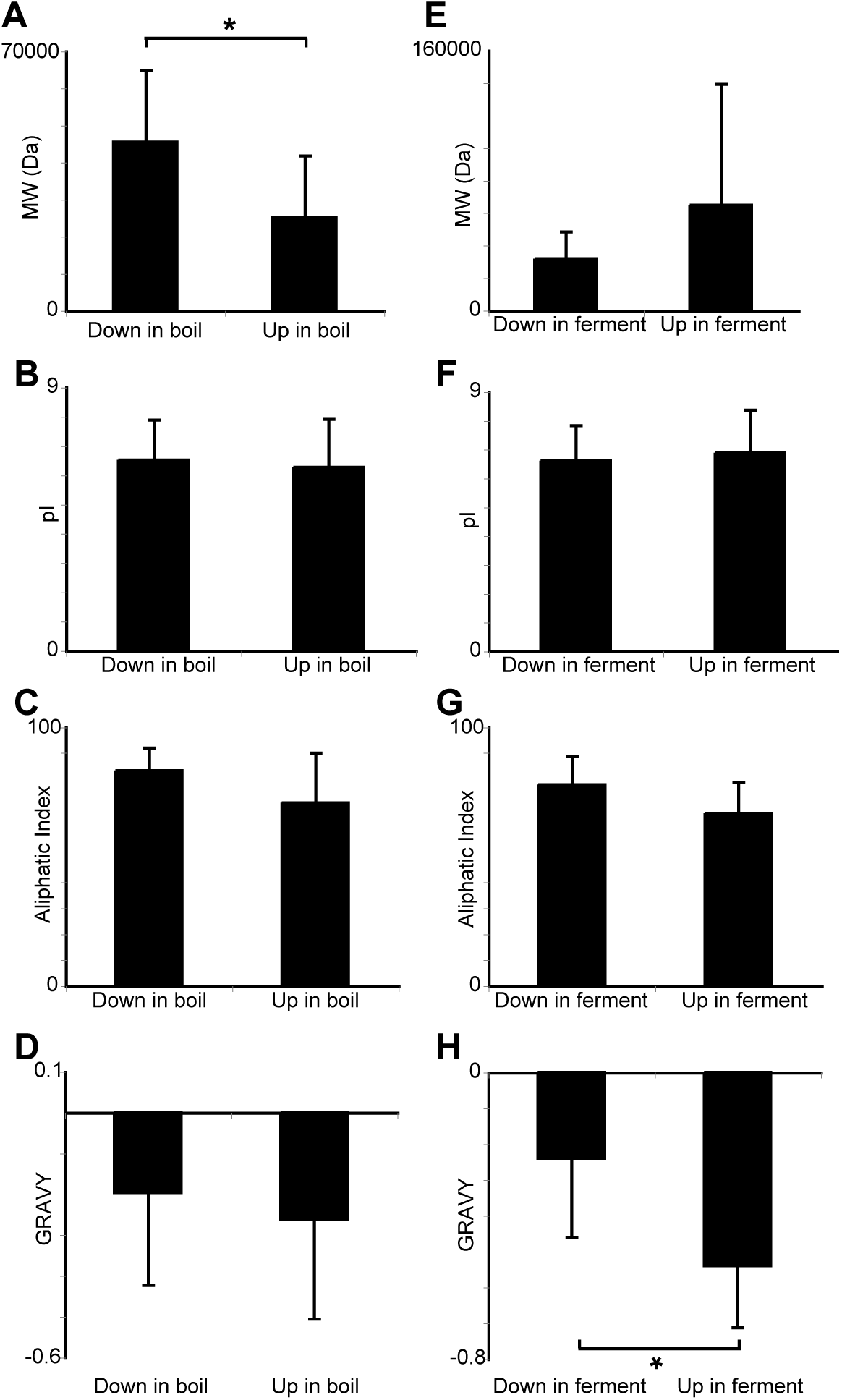
Biophysical properties of barley proteins changed in abundance from sweet wort to hopped wort, and to bright beer. (**A/E**) Molecular weight, (**B/F**) pI, (**C/G**) aliphatic index, and (**D/H**) grand average hydrophobicity (GRAVY) of proteins with large (log_2_ fold change > 1.5) and significant (P<0.05) changes in relative abundance as measured by SWATH between sweet wort and hopped wort (A,B,C,D), and between bright beer and hopped wort (E,F,G,H). Values are mean, n>10. Error bars are standard deviation. *, *P*<0.05 student’s T-test.

We next compared the effect of malting condition on the bright beer proteome. This comparison identified many diverse proteins with significantly altered relative abundances. Proteins with large and significant increases in relative abundance in high KI treatment relative to low KI treatment with coarse grist included thioredoxin-like protein, Gamma-hordein-3, Glutamate dehydrogenase, LTP2, B3-hordein, and B1-hordein (Fig. 5, Supplementary Table S2). The higher relative abundance of these proteins is consistent with more advanced germination with gibberellic acid treatment in the high KI samples allowing more efficient release and solubilization of these nitrogen storage proteins.

**Figure 5.**
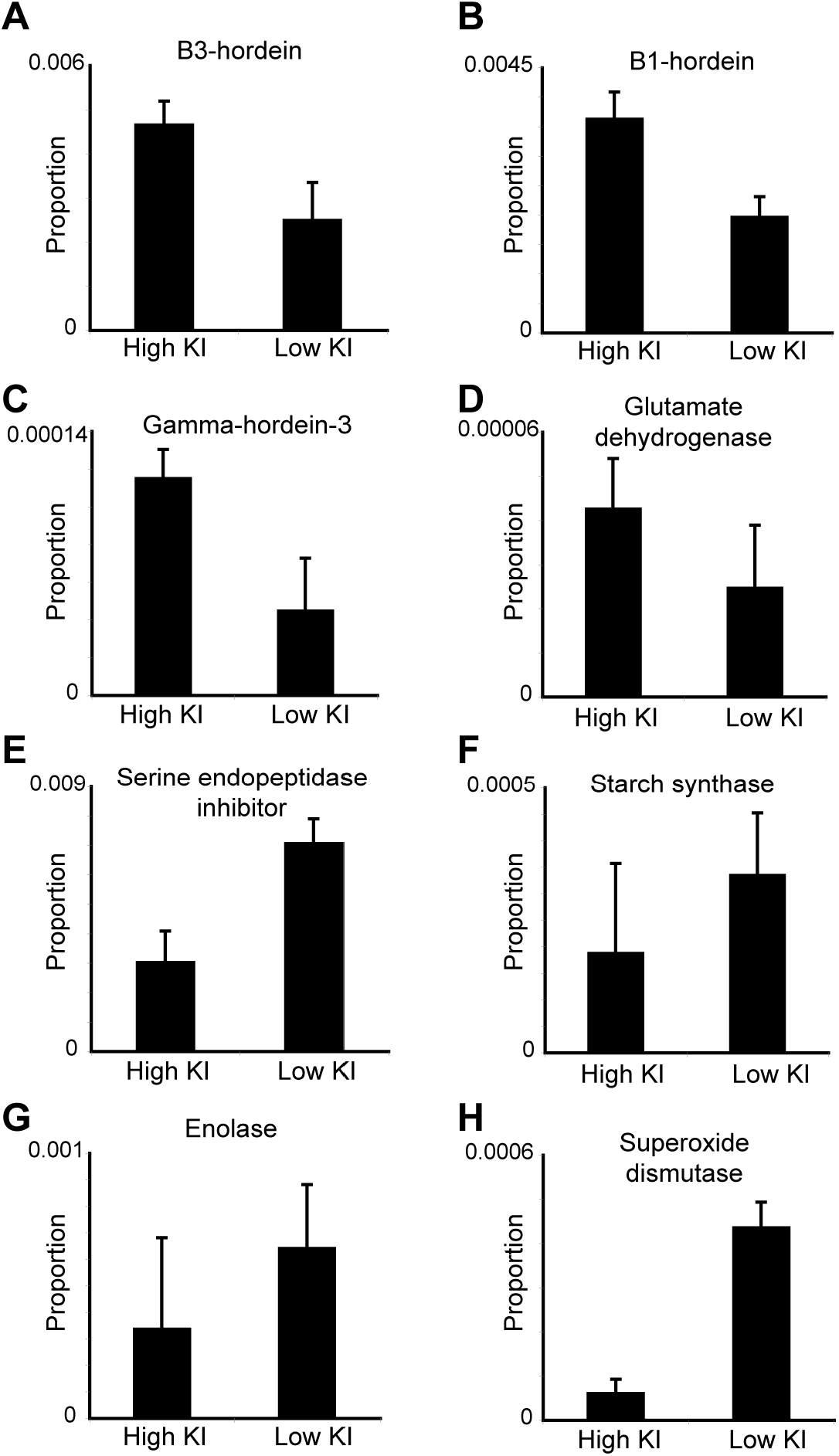
Malt modification changes the proteome of bright beer. Selected proteins with significant differences (P<0.05 for all comparisons) in relative abundance in bright beer made from high or low KI malt. (**A**) B3-hordein (P06471), (**B**) B1-hordein (P06470), (**C**) Gamma-hordein-3 (P80198), (**D**) Glutamate dehydrogenase (W5FCT5), (**E**) Serine endopeptidase inhibitor (M0XBS5), (**F**) Starch synthase (Q8H1Y8), (**G**) Enolase (F2D4W3), and (**H**) Superoxide dismutase [Cu-Zn] (Q96123). Values are mean, n=3. Error bars are standard deviation.

Chemical and enzymatic modification of proteins during malting, mashing, and fermentation is common, and includes oxidation, glycation, and proteolytic degradation. We examined our untargeted proteomic data to identify any such potential protein modifications. We identified numerous protein modifications, including glycation of lysines and oxidation of methionines, prolines, and tryptophans (Table 2). These modifications were often not stoichiometric, meaning that the final bright beer proteome was extremely diversely modified. Modifications such as glycation and oxidation are likely catalyzed by the heat and high concentrations of protein and reducing sugars present in wort, and are important contributors to the colour and flavour sensory properties of beer.

**Table 2.**
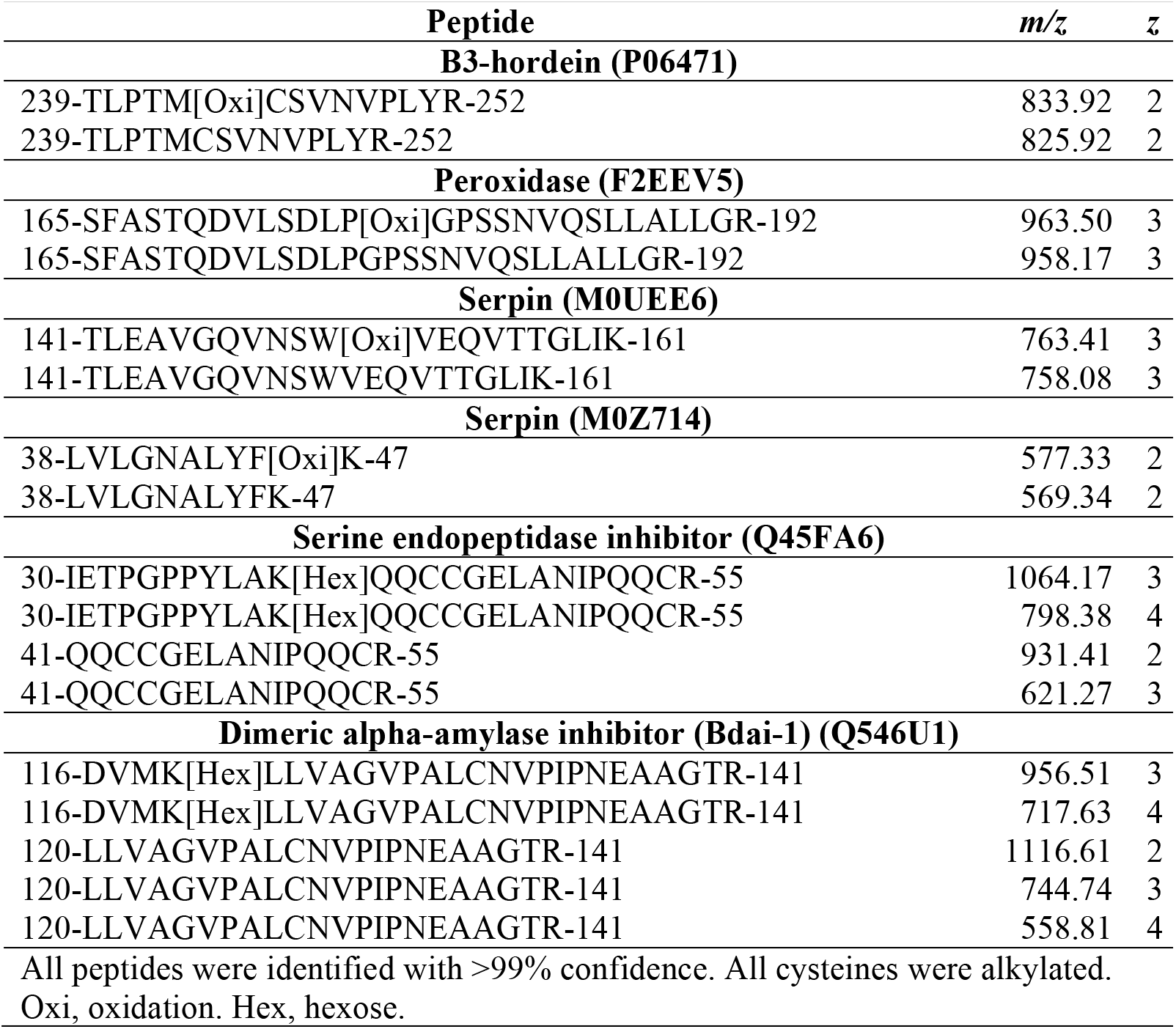
Selected covalently modified peptides and their unmodified forms from barley proteins identified in bright beer.

We identified many non-tryptic cleavage events in peptides (Table 3), which are most probably due to proteolytic cleavage from yeast or barley enzymes during malting, mashing, or fermentation. Altered malting conditions changed the relative abundance of several of these proteolytic cleavage events (Fig. 6A and B), consistent with the extent of germination during malting influencing protease expression, activity, and access to hordein, glutelin, and other protein substrates. Alignment of all sites of non-tryptic proteolysis in confidently identified peptides showed a clear preference for cleavage near prolines (Fig. 6C). This is likely because prolines are generally present at the ends of secondary structural elements, which are more likely to be surface exposed and accessible to proteases. The presence of these diverse proteolytic cleavage events further increased the complexity of the bright beer proteome, and would likely be influenced by the abundance of specific proteases, and the malting and mashing conditions used for beer production.

**Table 3.** Non-tryptic peptides from barley proteins identified in bright beer.

| Peptide | *m/z* | *z* |
| --- | --- | --- |
| **B1-hordein (P06470)** |  |  |
| 196-AIVYSIFLQ-204 | 527.30 | 2 |
| 196-AIVYSIFLQE-205 | 591.82 | 2 |
| 196-AIVYSIFLQEQ-206 | 655.85 | 2 |
| 196-AIVYSIFLQEQPQ-208 | 768.41 | 2 |
| 196-AIVYSIFLQEQPQQ-209 | 832.44 | 2 |
| 145-VFLQQQCSP-153 | 553.77 | 2 |
| 145-VFLQQQCSPV-154 | 603.30 | 2 |
| 145-VFLQQQCSPVP-155 | 651.83 | 2 |
| 145-VFLQQQCSPVPVP-157 | 749.89 | 2 |
| 145-VFLQQQCSPVPVPQR-159 | 891.97 | 2 |
| **B3-hordein (P06471)** |  |  |
| 167-AIVYSIVLQEQ-177 | 631.85 | 2 |
| 167-AIVYSIVLQEQS-178 | 675.37 | 2 |
| 167-AIVYSIVLQEQSL-179 | 731.91 | 2 |
| 167-AIVYSIVLQEQSLQ-180 | 795.94 | 2 |
| 167-AIVYSIVLQEQSLQL-181 | 852.48 | 2 |
| 239-TLPTMCSVN-247 | 511.74 | 2 |
| 239-TLPTMCSVNV-248 | 561.27 | 2 |
| 239-TLPTMCSVNVPL-250 | 666.34 | 2 |
| 239-TLPTMCSVNVPLY-251 | 747.87 | 2 |
| 239-TLPTMCSVNVPLYR-252 | 825.92 | 2 |
| **Gamma-hordein-3 (P80198)** |  |  |
| 144-EFLLQQCTL-152 | 576.29 | 2 |
| 144-EFLLQQCTLD-153 | 633.81 | 2 |
| 144-EFLLQQCTLDE-154 | 698.33 | 2 |
| 144-EFLLQQCTLDEK-155 | 762.37 | 2 |

All peptides were identified with >99% confidence. All cysteines were alkylated.

**Figure 6.**
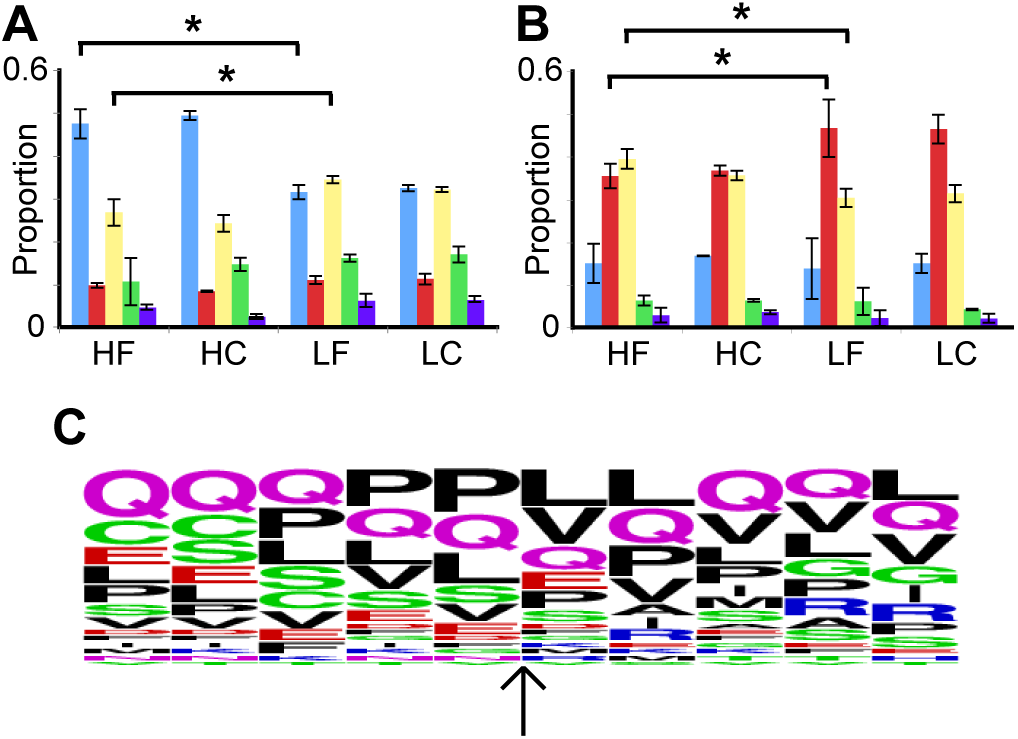
Non-tryptic proteolysis of barley proteins in bright beer. Relative abundance of different cleaved forms of peptides from B1-hordein (P06470) (**A**) _196_AIVYSIFLQ_204_ (blue), _196_AIVYSIFLQE_205_ (red), _196_AIVYSIFLQEQ_206_ (yellow), _196_AIVYSIFLQEQPQ_208_ (green), and _196_AIVYSIFLQEQPQQ_209_ (purple), and (**B**) _145_VFLQQQCSP_153_ (blue), _145_VFLQQQCSPV_154_ (red), _145_VFLQQQCSPVP_155_ (yellow), _145_VFLQQQCSPVPVP_157_ (green), and _145_VFLQQQCSPVPVPQR_159_ (purple). H, high KI; L, low KI; F, fine grist; C, coarse grist. Values are mean, n=3. Error bars are standard deviation. *, P<0.05 between HF and LF. These comparisons were also significant between HC and LC (not shown on graph). (**C**) WebLogo representation of aligned sequences from 31 non-tryptic cleavage sites detected in B1-hordein (P06470), B3-hordein (P06471), and Gamma-hordein-3 (P80198). Proteolysis occurred at the position of the arrow.

## Conclusions

The methods we present here for detailed analysis of the proteins and their modifications present at various stages throughout the beer production process demonstrate the opportunities offered by modern mass spectrometry proteomics in understanding the ancient process of beer production. We anticipate that these methods will be useful in detailed characterization of systems-level protein biochemistry in beer production and for improved quality control of the process.

## Acknowledgements

This research was supported by a University of Queensland Collaboration and Industry Engagement Fund grant to GPF, CEV, and ASN. BLS was supported by an NHMRC R.D. Wright Biomedical Career Development Fellowship Level 2 (APP1087975). CEV was supported by a Queensland Government Accelerate Fellowship. We acknowledge the technical assistance of The University of Queensland School of Chemistry and Molecular Biosciences Mass Spectrometry Facility.

